# Structural basis of nucleosome recognition by the human NuA4/TIP60 acetyltransferase complex

**DOI:** 10.64898/2026.11.03.763962

**Authors:** Zhenlin Yang, Alfredo Jose Florez Ariza, Jacques Côté, Eva Nogales

**Affiliations:** California Institute for Quantitative Biosciences (QB3), University of California, Berkeley, Berkeley, CA, USA; Howard Hughes Medical Institute, University of California, Berkeley, Berkeley, CA, USA; St-Patrick Research Group in Basic Oncology, Oncology Division of the CHU de Québec-Université Laval Research Center, Laval University Cancer Research Center, Quebec City, QC, Canada; Department of Molecular and Cell Biology, University of California, Berkeley, CA, USA; Molecular Biophysics and Integrative Bioimaging Division, Lawrence Berkeley National Laboratory, Berkeley, CA, USA

## Abstract

The human Tat-interactive protein, 60 kilodalton (Tip60) histone acetyltransferase (HAT) complex (NuA4/TIP60 complex, from now on TIP60-C) is a central regulator of chromatin accessibility, coupling histone acetylation with transcriptional activation and genome stability. But how the catalytic HAT module engages its nucleosome substrate has remained unknown. Here we present the structure of the human TIP60-C HAT module bound to the nucleosome, revealing a defined binding architecture in which TIP60, EPC1, ING3, and MEAF6 cooperatively position the catalytic center on the nucleosome surface, with the HAT module anchored on the histone acidic patch and engaging the linker DNA. The structure uncovers an intramolecular interaction involving the N-terminal Tudor domain of the TIP60 acetyltransferase subunit that suggests a potential mechanism for autoregulation of HAT activity. Together, our findings define the core principles by which the human TIP60-C HAT module recognizes chromatin, establishing a structural foundation for understanding how TIP60-C modifies chromatin in metazoans.

## INTRODUCTION

Acetylation of histone tails is a central mechanism for regulating chromatin structure and genome function^1–3^. In human cells, this process is mediated by multiple families of histone acetyltransferases (HATs) that often operate within large multisubunit complexes to integrate chromatin recognition with enzymatic activity^2^. These complexes regulate diverse nuclear activities, including transcription, DNA repair, replication, and genome maintenance^4,5^.

The TIP60 protein (also known as KAT5) belonging to the MYST family of acetyltransferases is part of the NuA4/TIP60 complex (TIP60-C), a ∼1.8-MDa assembly^6–11^ that acetylates both histone and non-histone substrates to regulate transcription, autophagy, DNA damage repair, and cell-cycle control^12–16^. TIP60-C combines activities that in yeast are distributed between two distinct complexes: the Nucleosome Acetyltransferase of histone H4 (NuA4) complex, which catalyzes histone acetylation^8,17,18^, and the switch/sucrose nonfermentable related 1 (SWR1) complex, which mediates H2A.Z histone variant deposition^19,20^. This functional convergence suggests that human TIP60-C may engage nucleosomes in ways that differ from yeast NuA4.

Biochemical and structural analysis of the yeast NuA4-HAT module (piccolo NuA4 or picNuA4), comprising Esa1 (the catalytic subunit), Epl1, Yng2, and Eaf6, has shown that it binds both nucleosomal DNA and the core histones, recognizing the H2A-H2B acidic patch and positioning the catalytic Esa1 subunit near the histone H4 N-terminal tail^17,21–24^. In this arrangement, the linker DNA is oriented toward the large Tra1 subunit and does not directly engage the HAT module^23,24^.

The human TIP60-C HAT module consists of TIP60, EPC1, ING3, and MEAF6^17^ and contains conserved chromatin-interacting domains, including the PHD finger of ING3 that recognizes histone H3 tail methylated on lysine 4 ^17,25,26^, and the N-terminal Tudor domain of TIP60, which has been proposed to interact with modified histone peptides^27–29^. Recent studies suggest that the TIP60 Tudor domain can influence global H4 acetylation levels through mechanisms independent of chromatin recruitment^30^, like its yeast homolog^31^, raising the possibility that it participates in regulatory interactions within the HAT module rather than serving solely as a chromatin-targeting element.

Here, we used cryo-EM to obtain the structure of the nucleosome-bound human TIP60-C HAT module. The structure, together with complementary biochemical analyses, defines the molecular basis of nucleosome engagement and reveals features that differ from those seen for the yeast NuA4 complex. Our work provides insight into how multi-subunit HAT complexes interact with nucleosomal substrates and regulate histone acetylation, and suggests how the HAT module-mediated interactions may contribute to chromatin engagement by the intact TIP60-C complex^32^.

## RESULTS

### Overall structure of the TIP60-C HAT module bound to the nucleosome

To define how human TIP60-C engages its nucleosomal substrate, we reconstituted the HAT module with full-length TIP60, ING3, MEAF6, and a truncated EPC1 (residues 1-350) (Fig. 1a) and carried out structural analysis of the module bound to a nucleosome (Extended Data Fig. 1-3). The purified complex was catalytically active, acetylating histone H4 in an acetyl-CoA–dependent manner (Fig. 1b). For structural analysis, the active HAT module was incubated with nucleosomes bearing a 22 bp of linker DNA and reconstituted using histone octamers containing a carboxymethyl coenzyme A (CMC)-modified histone H4K16 to maximize binding ^23^ (Extended Data Fig. 1a,b).

**Figure 1.**
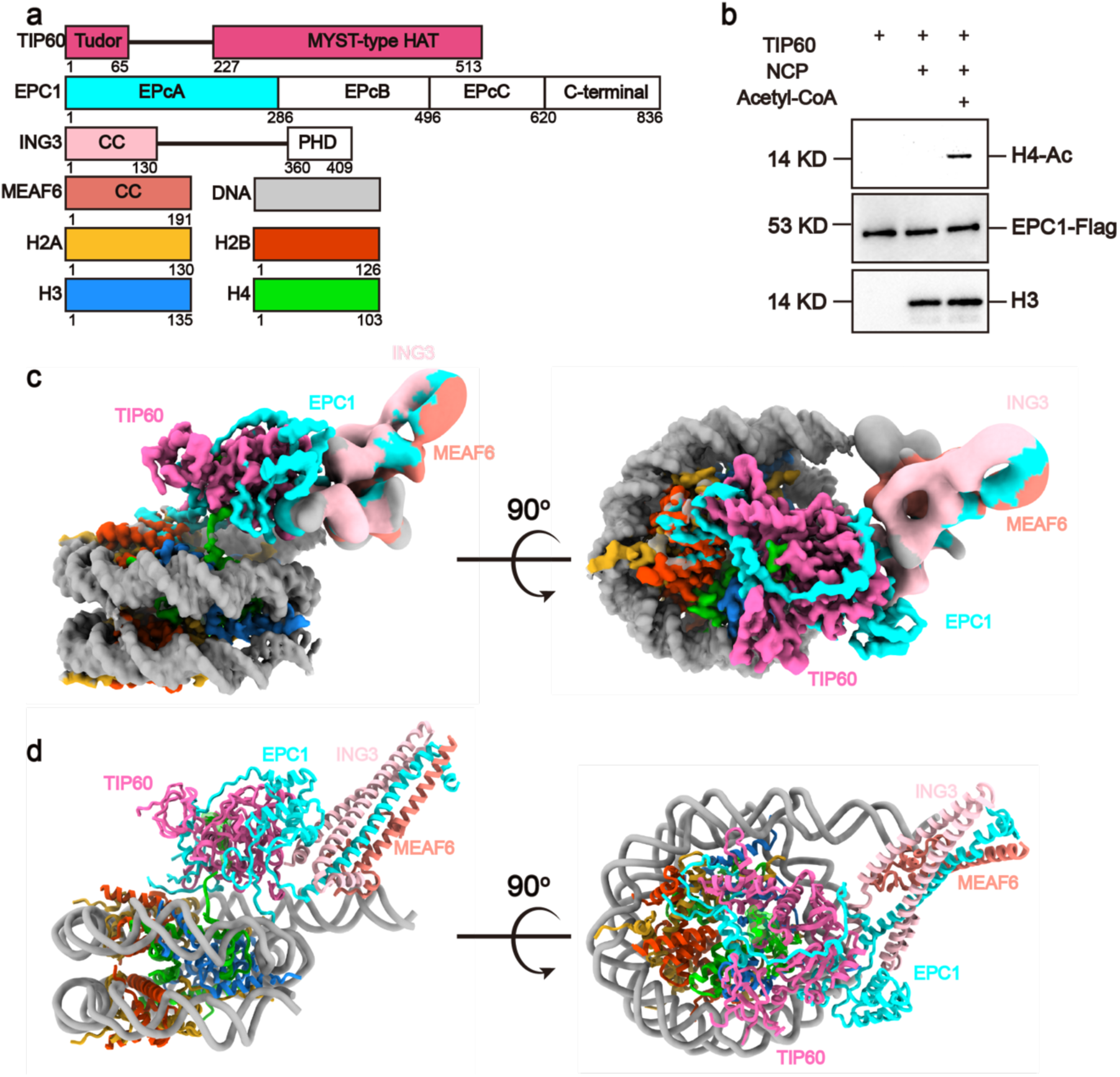
Structure of the human TIP60 HAT module bound to a nucleosome. **a**, Domain organization of the subunits comprising the human TIP60-C HAT module. The color scheme is used consistently throughout the figures. **b**, In vitro histone acetyltransferase (HAT) assay showing that the purified TIP60-C HAT module is catalytically active on nucleosomal substrates, acetylating histone H4 in an acetyl-CoA–dependent manner. **c**,**d**, Two orthogonal views of the cryo-EM density map (**c**) and atomic model (**d**). The HAT module adopts a clamp-like architecture that engages the nucleosome surface, with EPC1, ING3, and MEAF6 contributing to position the catalytic TIP60 subunit adjacent to the H4 tail exit site.

We used single-particle cryo-EM of the TIP60-C HAT-nucleosome complex to produce a reconstruction with overall resolution of 2.5 Å and 2.9 Å for the catalytic core of the HAT module, respectively, (Fig. 1c,d; Extended Data Figs. 2,4,5), enabling atomic modeling. The structure shows the TIP60 catalytic center engaged with the CMC-modified H4 N-terminal tail region, consistent with its substrate preference (Fig. 1d). EPC1, ING3, and MEAF6 form an extensive interaction surface with the nucleosome (Fig. 1c,d), including engagement with the H2A-H2B acidic patch, which is well resolved in the map (Fig. 1c,d). Additionally, the structure suggests a potential interaction with linker DNA, as one end of the nucleosomal DNA locates adjacent to a helical bundle formed by EPC1, ING3, and MEAF6 (Fig. 1c,d; Extended Data Fig. 3).

### Acidic patch recognition by EPC1 and TIP60 anchors the complex to the nucleosome

A prominent interface between the TIP60-C HAT module and the nucleosome is centered on the H2A-H2B acidic patch (Fig. 2a-c). EPC1 contributes two conserved arginine residues, Arg7 and Arg36, which insert into the acidic pocket formed primarily by H2A Glu61, Glu64, and Glu92 to establish salt-bridge and hydrogen-bond interactions (Fig. 2b). The arginines are located within the N-terminal EPcA region of EPC1 (Fig.1a), a segment enriched in positively charged amino acids.

**Figure 2.**
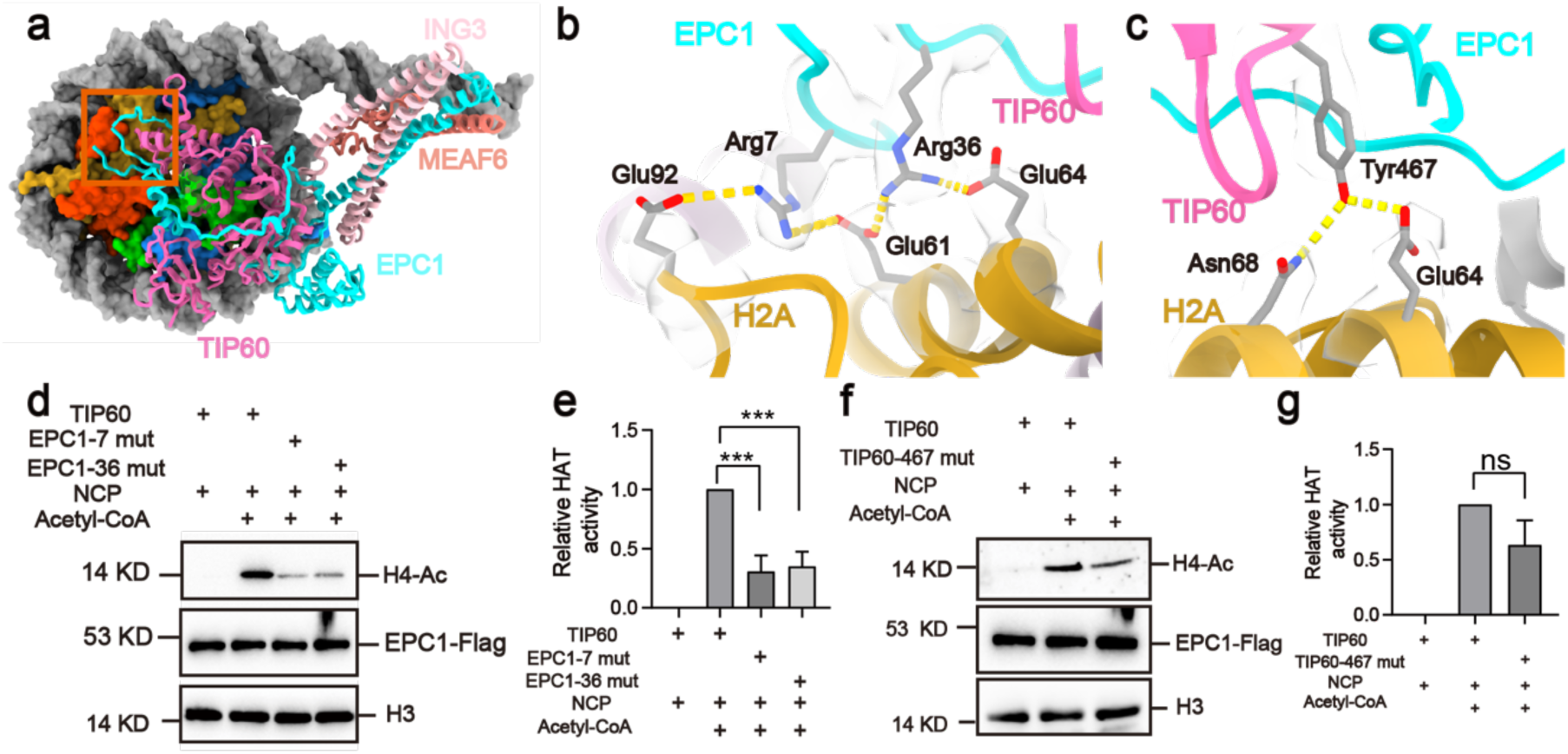
Anchoring of the TIP60-C HAT module to the nucleosome acidic patch. **a**, Top view of the TIP60-C HAT-nucleosome structure. The red box marks the region of HAT interaction with the H2A–H2B acidic patch highlighted in the following panels. **b**, Details of the ‘arginine-anchor” mechanism with the acidic patch, in which EPC1 residues Arg7 and Arg36 interact with H2A residues Glu61, Glu64, and Glu92. **c**, Additional contacts between the Tip60 C-terminal region and the nucleosome include a hydrogen bond between Tip60 Tyr467 and H2A Glu64. **d**, Western blot of HAT activity assays quantifying nucleosomal H4 acetylation by HAT complexes containing EPC1 wild-type, and R7A or R36A mutants. **e**, Quantitation of d showing mean ± s.d., (n = 3 independent experiments, *** P < 0.001). **f**,**g**, As **d**,**e** comparing HAT complex containing wild type and Y467A mutant TIP60.

In yeast, the EPC1 homolog Epl1 engages the acidic patch through Arg56 and Arg58^23^ (Extended Data Fig. 6), corresponding to human Arg7 and Arg9. Despite the conservation, EPC1 Arg9 does not contact the acidic patch, which instead is engaged by the sequence-distant Arg36, giving the EPC1 N-terminal region a distinct anchoring geometry from its yeast counterpart.

In addition to EPC1, the C-terminal segment of the catalytic subunit TIP60 approaches the H2A-H2B dimer near the acidic patch, with Tyr467 hydrogen-bonding with H2A Glu64 (Fig. 2c). This placement brings the TIP60 C-terminus near the acidic patch and contributes to a composite interface between the HAT module and the nucleosome core.

We assessed the functional importance of EPC1-mediated acidic-patch engagement by mutational analysis. Both the EPC1 Arg7Ala and the Arg36Ala mutants exhibited a strong reduction in H4 acetylation relative to the wild-type complex (Fig. 2d,e). In contrast, mutation of TIP60 Tyr467 to alanine did not result in a statistically significant reduction in HAT activity under the conditions tested (Fig. 2f,g). These results support an important role for EPC1-mediated acidic-patch recognition in efficient nucleosome acetylation by the TIP60-C HAT module.

### Linker DNA engagement by MEAF6

The TIP60-C HAT-nucleosome structure reveals additional density extending from the HAT core towards the exiting linker DNA. Although this region is flexible and the resolution is lower than for the rest of the complex, we could assign to a helical bundle formed by portions of ING3, MEAF6, and EPC1 with the aid of AlphaFold3 structure prediction (Fig. 3a; Extended Data Fig. 3). The fitting of the AlphaFold3 model into the density map suggests that an alpha helix in MEAF6 sits by the nucleosomal DNA entry/exit site, with the conserved Lys74, Arg78, and Arg80 residues forming a cluster that engages the linker DNA (Fig. 3b). Sequence alignment of MEAF6 across species shows that these residues are conserved in metazoans but are not conserved in the corresponding region of yeast Eaf6 (Fig. 3c). This conservation pattern is in agreement with the fact that linker DNA engagement is not observed for the homologous Eaf6 subunit in the yeast NuA4 nucleosome structure^23,24^, highlighting a divergence in extranucleosomal DNA interactions between the human and yeast complexes.

**Figure 3.**
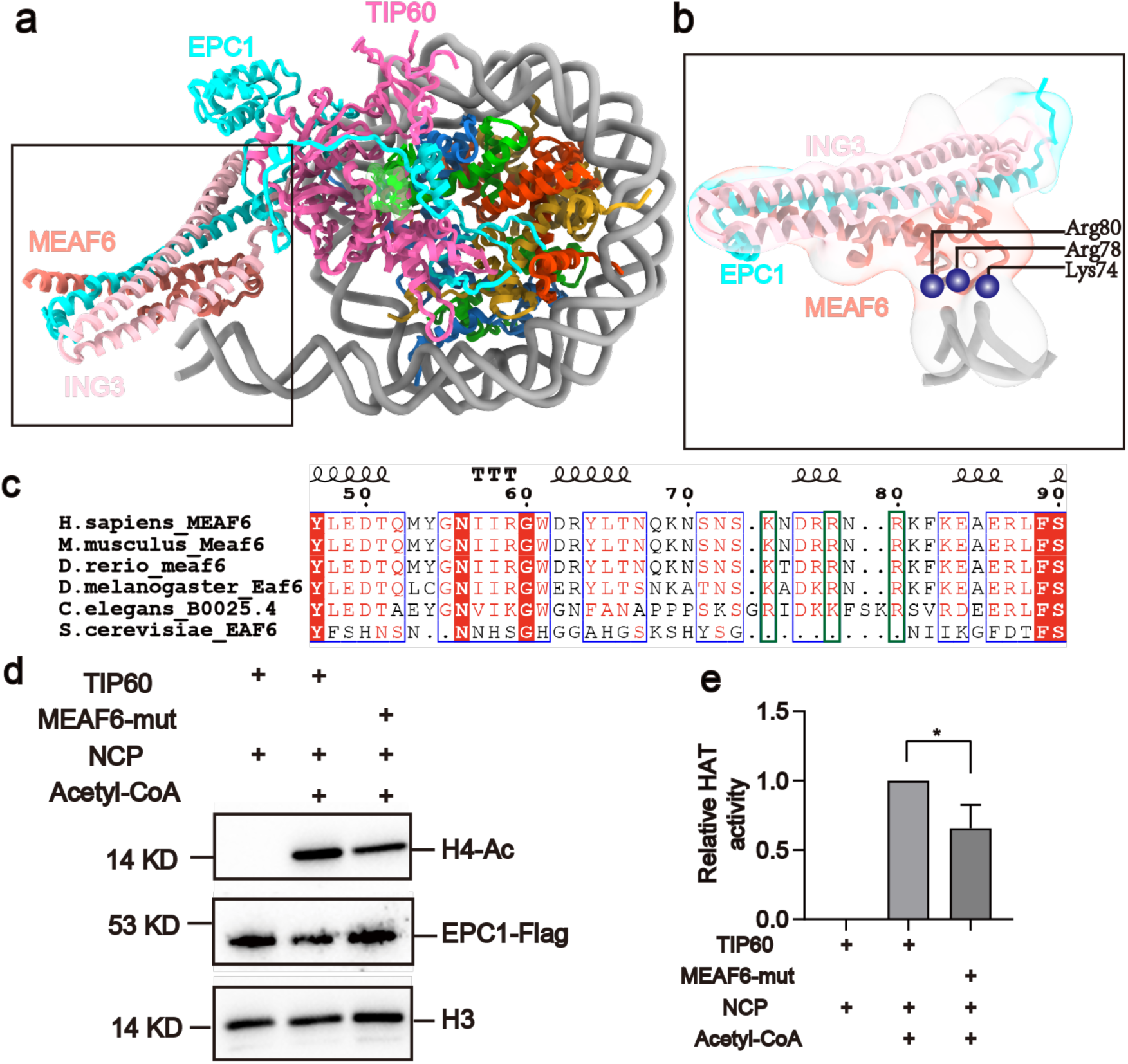
Putative MEAF6-mediated binding to linker DNA. **a**, Overall structure of the human TIP60-C HAT module bound to the nucleosome. The region of putative linker DNA interaction is highlighted by the black-outlined box, and shown in more detail in b. **b**, Close-up view of the helical bundle formed by EPC1, ING3, and MEAF6, showing the positioning of MEAF6 near the linker DNA. **c**, Sequence alignment across different species generated using ESPript (REF?). Red blocks highlight totally conserved residues. Green boxes indicate residues proposed to mediate linker DNA interactions. **d**-**e**, Western blot (left) and quantitation (right; mean ± s.d., n = 3, * P < 0.05) of H4 acetylation by the TIP60-C HAT incorporating either wild type or mutant MEAF6 with K74D, R78D and R80D.

To assess the functional importance of this interface, we generated a MEAF6 triple-Aspartate mutant (K74D/R78D/R80D) and measured its HAT activity. The mutant exhibited a moderate but statistically significant decrease in H4 acetylation relative to the wild-type complex (Fig. 3d,e). These results suggest that the basic surface of MEAF6 contributes to nucleosome acetylation, potentially by stabilizing the interaction between the TIP60-C HAT module and the chromatin substrates.

### The TIP60 Tudor domain contacts the HAT catalytic site

Our structure reveals an intramolecular interaction of the TIP60 N-terminal Tudor domain with the catalytic pocket of the MYST-type domain within the protein (Fig. 4a). A region within a flexible loop in the Tudor domain (residues ∼50–60) extends toward the active site groove, an interaction that is not present in the yeast complex^23^, where the Tudor domain is not visible due to flexibility.

**Figure 4.**
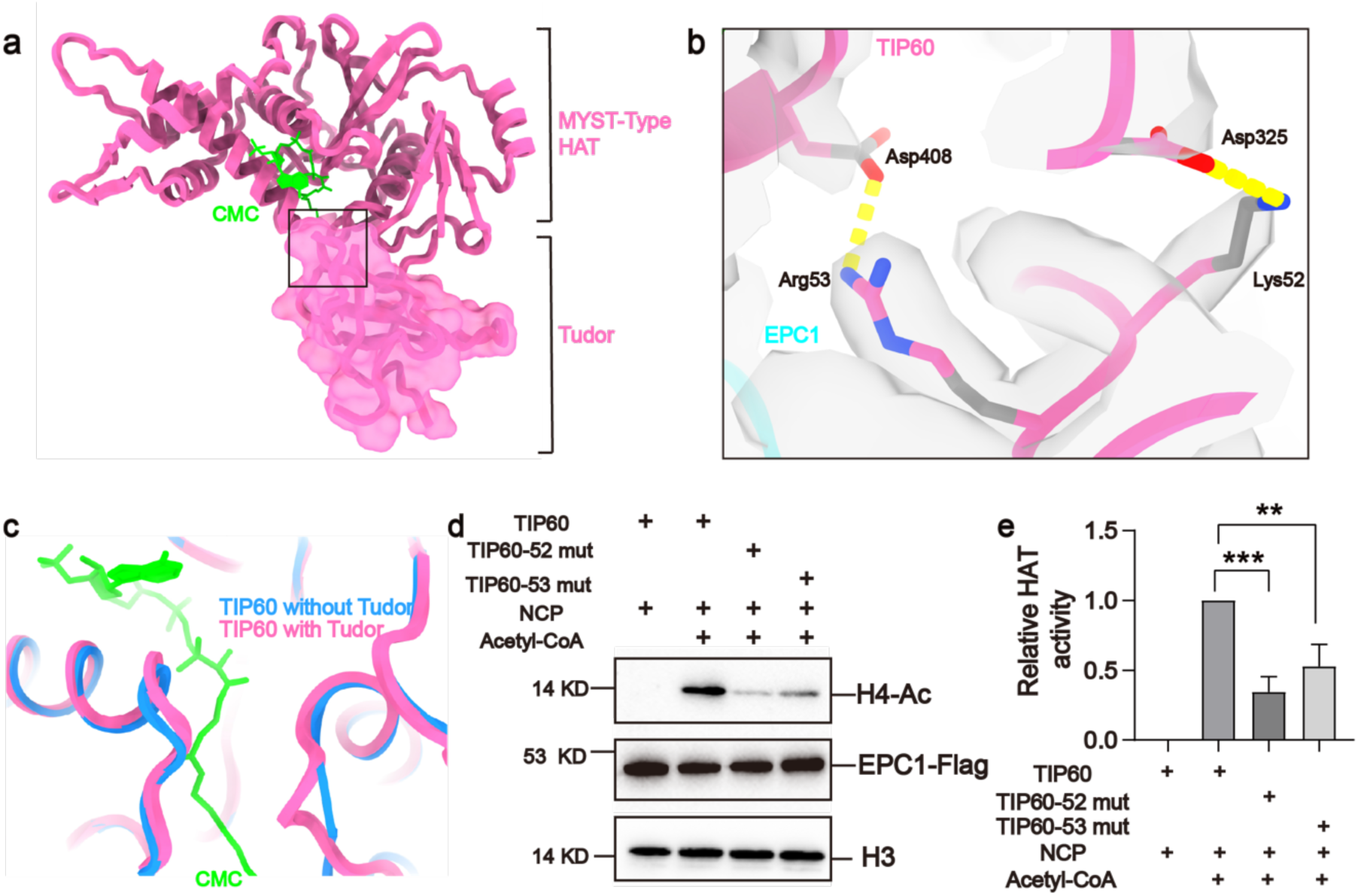
Cis regulation within the TIP60 catalytic subunit. **a**, Overview of the intramolecular interaction within the TIP60 catalytic subunit. The N-terminal Tudor domain (shown as a transparent surface), folds back onto the MYST catalytic core (highlighted by the black-outlined box and shown in **b)**. **b**, Details of the intramolecular interface within TIP60. A flexible loop (residues 50–60) extends towards the active-site groove, where Lys52 and Arg53 form stabilizing electrostatic interactions with Asp325 and Asp408 in the catalytic core, adjacent to the acetyl-CoA binding pocket. **c**, Structural comparison of the TIP60 catalytic core in the presence (pink, this work) and absence of the Tudor domain (light blue, PDB: 2OU2). The CMC moiety is shown in green. **d**,**e**, HAT activity assays for TIP60-C HAT variants, comparing wild type with TIP60 K52A and R53A mutants disrupting this intramolecular contact. (**d**, western blot) (**e**, quantification showing mean ± s.d., n = 3, ** P < 0.01, *** P < 0.001).

TIP60 residues Lys52 and Arg53 in the Tudor domain of the human protein contact Asp325 and Asp408 within the MYST catalytic core, forming electrostatic interactions that anchor the Tudor loop near the catalytic center (Fig. 4b). Sequence alignment shows that Lys52 and Arg53 are conserved across metazoan species (Extended Data Fig. 7), underscoring the functional importance of this intramolecular interface. Furthermore, comparison of our HAT module structure with the crystal structure of the isolated Tip60 MYST catalytic core lacking the Tudor domain (PDB: 2OU2) reveals that the presence of the Tudor domain is associated with a more closed conformation of the acetyl-CoA binding pocket, suggesting that these intramolecular interactions may influence the acetyl-CoA–binding pocket (Fig. 4c). To assess the functional importance of this interface, we introduced alanine substitutions at TIP60 Lys52 and Arg53 and measured the histone acetylation activity by the reconstituted TIP60-C HAT module. Mutation of either residue resulted in a marked reduction in HAT activity, consistent with our previous finding that mutation of Arg53 also affects the activity of the native TIP60-C^33^. These results support a role for Tudor-mediated engagement of the catalytic core in maintaining HAT catalytic activity (Fig. 4d,e).

## DISCUSSION

Our cryo-EM structure of the human nucleosome-bound TIP60-C HAT module reveals a clamp-like recognition mechanism, in which the enzyme engages its nucleosomal substrate through a bipartite interaction with both the histone acidic patch and the linker DNA for efficient H4 acetylation. The structure also points to key evolutionary adaptations that distinguish the human TIP60 complex from its yeast counterparts (Fig. 5), likely reflecting the increased complexity of the metazoan epigenome.

**Figure 5.**
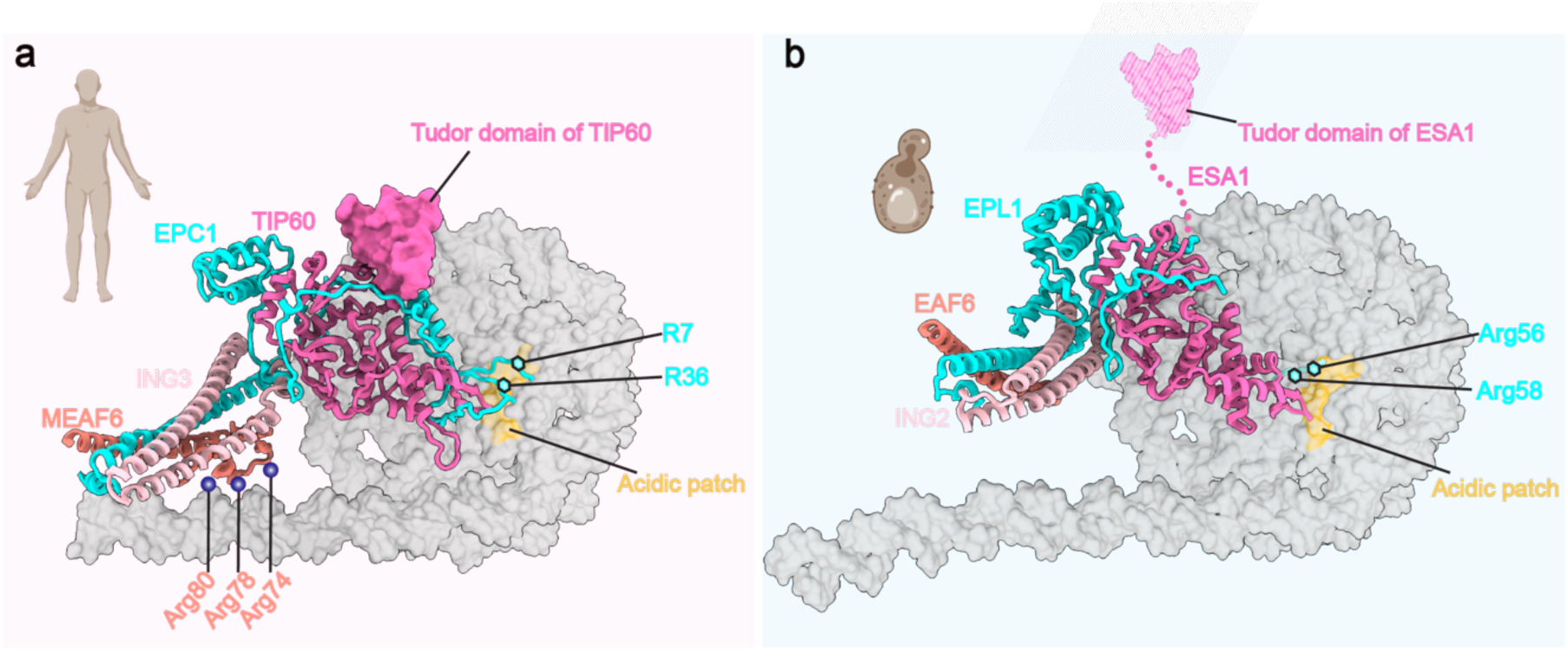
Structural comparison of nucleosome recognition by human TIP60-C and yeast NuA4 HAT modules. **a**, Structure of the human TIP60 HAT module bound to the nucleosome, highlighting acidic-patch recognition by EPC1 Arg7 and Arg36, linker DNA engagement by MEAF6, and the intramolecular interaction between the TIP60 Tudor domain and its catalytic core. **b**, Structure of the yeast NuA4 HAT module bound to the nucleosome, showing acidic-patch recognition by Epl1 Arg56 and Arg58. Unlike human MEAF6, yeast Eaf6 does not engage linker DNA, and the Esa1 Tudor domain is unresolved due to its flexibility.

A central feature of our structure is the anchoring of TIP60-C to the nucleosome through the EPC1 subunit. Targeting of the acidic patch is a strategy shared by diverse chromatin-acting factors, including BRCA1, PRC1, RCC1, and the yeast NuA4 complex^23,24,34–39^. Unexpectedly, in the human TIP60-C, EPC1 engages the acidic patch using two arginine residues that are distant in the sequence, Arg7 and Arg36 (Fig. 2b; 5a), rather than the compact Arg56/Arg58 motif observed in the homologous yeast Epl1^23^ (Fig. 5b). Given that the acidic patch can be altered by histone variants such as H2A.Z and by post-translational modifications including H2BK120ub, this altered mode of acidic-patch engagement may contribute to context-dependent interactions of TIP60 with chromatin.

Our structure also reveals density consistent with an interaction between the MEAF6 subunit and linker DNA, a feature not observed for the homologous Eaf6 subunit in yeast NuA4^23,24^ (Fig. 3a, b, d), suggesting an additional mode of nucleosome engagement by the human TIP60-C HAT module that may correlate with the increased prevalence of longer linker DNA in metazoan chromatin.

Finally, the structure provides insight into the role of the TIP60 Tudor domain, which has been interpreted to act either as a methylated histone reader or as a regulatory element required for HAT activity^30^ (Fig. 4d,e). Our structure and mutational analysis now support a model in which intramolecular contacts between the Tudor domain and the catalytic core help maintain HAT activity. Previous studies showed that the yeast Tudor domain is required for HAT activity on nucleosomes but not on free histones^40^, suggesting a role in nucleosome engagement. However, the Tudor domain has not been resolved in any picNuA4-nucleosome structure, nor in human MSL–nucleosome structures^22–24,41^, with only the structure of the Drosophila MSL showing a stable Tudor domain engaged with the nucleosome^42^. These observations indicate that Tudor domain interactions with nucleosomes may differ among chromatin-modifying complexes. Our structure reveals how the human TIP60 Tudor domain contacts the catalytic core, providing a structural basis for its role in regulating HAT activity in the human complex.

Collectively, our findings point to a modular evolution of the TIP60-C HAT subcomplex that parallels the large reorganization of the human TIP60-C with respect to its yeast counterparts NuA4 and SWR1. Although the catalytic core in TIP60 is conserved, peripheral attachment modules in the HAT module, such as EPC1 and MEAF6, appear to have adapted to the more nuance features of human chromatin, with EPC1 sensing the nucleosomal acidic patch and MEAF6 engaging linker DNA (Fig5). This structural framework clarifies the mechanism of TIP60-C-mediated acetylation and provides the basis for the rational design of inhibitors targeting interfaces unique to the TIP60-C HAT module, with potential relevance to cancers associated with TIP60 dysregulation.

## METHODS

### TIP60-C HAT expression and purification

The cDNAs encoding full-length human TIP60, ING3, and MEAF6 were individually cloned into the pCAG expression vector without affinity tags. A truncated form of EPC1 (residues 1–350) was cloned into the same vector with an N-terminal 3×FLAG tag. Equal amounts of the four plasmids (1.5 mg total DNA) were co-transfected into Expi293 cells using PEI 25k (Polysciences) following the manufacturer’s protocol. Cells were maintained at 37 °C in 8% CO₂ and harvested 72 h after transfection.

Cell pellets were resuspended in lysis buffer containing 150 mM NaCl, 20 mM HEPES (pH 8.0), 0.1% NP-40, protease inhibitor cocktail, 1 mM TCEP, and 10% (v/v) glycerol, and lysed by sonication. Insoluble material was removed by centrifugation at 40,000 rpm for 30 min at 4 °C. The clarified lysate was incubated with anti-FLAG M2 affinity resin for 3 h at 4 °C with gentle rotation. After three washes with lysis buffer, bound proteins were eluted with buffer supplemented with 0.5 mg mL⁻¹ 3×FLAG peptide. The eluate was further purified by anion-exchange chromatography on a 1 mL HiTrap Q column (Cytiva) using a linear NaCl gradient from 150 mM to 1 M. Fractions containing the TIP60-C HAT module were pooled and subjected to size-exclusion chromatography on a Superose 6 Increase 10/300 GL column (Cytiva) equilibrated in buffer containing 150 mM NaCl, 20 mM HEPES (pH 8.0), 0.01% NP-40, protease inhibitor cocktail, 1 mM TCEP, and 1% (v/v) glycerol. Peak fractions were pooled, concentrated, flash-frozen in liquid nitrogen, and stored at −80 °C for subsequent biochemical and structural analyses.

### Nucleosome reconstitution

The 147 bp Widom 601 DNA sequence containing an additional 22 bp linker was amplified by PCR and purified using a HiTrap Q column (Cytiva). Recombinant human histones H2A, H2B, and H3 were expressed in *E. coli* BL21(DE3) and purified from inclusion bodies as described previously. CMC-modified human histone H4 was generated by native chemical ligation following established protocols^23^.

Equimolar amounts of H2A, H2B, H3, and CMC-modified H4 were combined to a final concentration of 1 mg mL⁻¹ and denatured in buffer containing 6 M guanidine hydrochloride. The histone mixture was subsequently dialyzed into refolding buffer containing 2 M NaCl, 20 mM Tris-HCl (pH 7.5), 1 mM EDTA, and 5 mM β-mercaptoethanol. Refolded histone octamers were purified by size-exclusion chromatography on a Superdex 200 Increase 10/300 GL column (Cytiva). Purified histone octamers were mixed with DNA at a 1:0.9 molar ratio and assembled into nucleosomes by stepwise salt-gradient dialysis from 2 M to 10 mM NaCl. Nucleosome assembly was evaluated by native PAGE and visualized using SYBR Safe staining (Ext. Data Fig. 1a, lane2). Fractions containing nucleosomes were pooled and concentrated to approximately 1 µM.

### Electrophoretic mobility shift assays (EMSA)

EMSAs were performed to assess the binding of the TIP60-C HAT module to CMC-modified nucleosomes. CMC-modified nucleosomes, at a final concentration of approximately 100 nM, were incubated with increasing concentrations of the purified TIP60-C HAT module at the indicated HAT molar ratios. The samples were incubated at 4 °C for 30 min in a buffer containing 150 mM NaCl, 20 mM HEPES (pH 8.0), and 1 mM TCEP. Protein-nucleosome complexes were resolved by native PAGE using a 5% polyacrylamide gel and visualized by SYBR Safe staining (Ext. Data Fig. 2a).

### HAT activity assay

For in vitro acetylation assays, 15 nM of native or mutant TIP60-C HAT complex was incubated with 100 nM nucleosomes in the presence of 50 µM acetyl-CoA in reaction buffer containing 20 mM HEPES (pH 8.0), 100 mM NaCl, 0.1 mg/mL BSA, and 10 mM sodium butyrate. Reactions were carried out at 37 °C for 60 min and terminated by the addition of SDS loading buffer followed by heating at 95 °C for 1 min. Samples were analyzed by SDS–PAGE and western blotting using an anti-H4K5ac antibody (Thermo Fisher Scientific). Signal intensities were quantified using ImageJ. All assays were independently repeated three times.

### HAT-Nucleosome complex assembly and negative-Stain EM

CMC-modified nucleosomes and TIP60-C HAT complex were mixed at a 1:4 molar ratio and incubated on ice for 30 min. The mixture was loaded onto a glycerol gradient prepared with a top buffer containing 150 mM NaCl, 20 mM HEPES (pH 8.0), 1 mM TCEP, and 10% (v/v) glycerol, and a bottom buffer containing 150 mM NaCl, 20 mM HEPES (pH 8.0), 1 mM TCEP, 30% (v/v) glycerol, and 0.02% glutaraldehyde^43^. Gradient centrifugation was performed at 40,000 rpm for 14 h in a Beckman MSL50 rotor. Fractions were collected and analyzed by 5% native PAGE to assess complex formation and stability (Ext. Data Fig. 1B). The fractions enriched in the TIP60-C HAT-nucleosome complex were pooled and dialyzed against buffer containing 150 mM NaCl, 20 mM HEPES (pH 8.0), and 1 mM TCEP to remove glycerol.

The HAT–NCP complex was adjusted to a final concentration of approximately 100 nM and used for both negative-stain electron microscopy and cryo-EM grid preparation. For negative staining, 3.5 µL of sample was applied to glow-discharged continuous carbon grids and incubated for 5s. Excess liquid was blotted with filter paper, followed by three washes with 2% (w/v) uranyl formate. Grids were air-dried and subsequently examined using a Tecnai T12 transmission electron microscope (Thermo Fisher Scientific) operated at 120 keV. Images were acquired at a nominal magnification of 43,000× using a TVIPS camera (Ext. Data Fig. 2A).

### Cryo-EM grid preparation and data collection

For cryo-EM, 4 µL sample was applied to glow-discharged homemade graphene oxide coated UltrAuFoil R1.2/1.3 200-mesh gold grids^44^ and incubated at 4 °C for 2 min. Grids were blotted with a blot force of 5 and blot time of 3.5 s, and plunge-frozen into liquid ethane using a Vitrobot Mark IV (Thermo Fisher Scientific).

Grids were initially screened using a Talos Arctica electron microscope (Thermo Fisher Scientific). Suitable grids were transferred to a Titan Krios G3 (Thermo Fisher Scientific) at the Cal-Cryo facility for data collection. Movies were recorded using SerialEM at a calibrated pixel size of 1.05 Å^45^. Each movie was collected with a total dose of 50 e⁻/Å² fractionated into 50 frames. A total of 13,322 movies were acquired (Ext. Data Fig. 2B).

### Cryo-EM Data Processing

Cryo-EM movies were imported into CryoSPARC and subjected to Patch Motion Correction followed by Patch CTF Estimation^46^. Initial particles were automatically picked from 3,000 micrographs using the Blob Picker, followed by several rounds of 2D classification. 2D class averages exhibiting extra density corresponding to the HAT module were selected as templates for template-based particle picking on the full dataset, yielding a total of 8,682,166 particles. These particles were initially extracted with a box size of 80 pixels at a pixel size of 4.2 Å and split into two independent datasets. Each dataset underwent two rounds of 2D classification, and particles from classes displaying clear secondary structural features were retained (Ext. Data Fig. 2C).

The selected particles were re-extracted with a box size of 320 pixels at the original pixel size of 1.05 Å, yielding a final dataset of 1,955,409 particles. These particles were imported into RELION for further processing⁴⁰. An initial 3D auto-refinement was performed using the CryoSPARC ab initio map as a reference. Subsequently, focused 3D classification with a soft mask around the HAT catalytic core was carried out to isolate the most stable class exhibiting consistent and well-resolved HAT core density. The best-resolved class (harboring 394,258 particles) was imported into CryoSPARC for non-uniform refinement, followed by local refinement using a mask focused on the HAT catalytic core, yielding a high-resolution density map of the catalytic core. This map was used for atomic model building and structural refinement.

To enhance the signal corresponding to the helical bundle region of the HAT module, starting from the 3D refined structure using the 1,955,409 particles imported into RELION, we designed a soft mask encompassing the nucleosome region and inverted it, generating an “inverted mask” corresponding to a cubic volume containing a cavity that matches the well-resolved nucleosome density (Ext. Data Fig. 3). This mask was used for particle signal subtraction, followed by re-centering on the nucleosome center of mass and re-boxing to a final box size of 220 × 220 pixels. The resulting signal-subtracted particles were subjected to 2D classification without alignment. Several 2D class averages displaying clear features corresponding to the HAT module were retained, whereas classes lacking discernible structural features were discarded.

A total of 159,808 particles were selected and 3D back-projected, with the resolution limited to 10 Å, to generate a low-pass filtered reference for subsequent 3D auto-refinement. The resulting density map revealed the HAT module in its full spatial extent, comprising the catalytic core (hereafter referred to as body 1) and the helical bundle region (hereafter referred to as body 2). Masks corresponding to each body were generated and used for an initial round of multi-body refinement. This analysis recovered signal for both bodies, albeit at limited resolution. Analysis of the multi-body eigen-vector/eigen-value results revealed a unimodal distribution of amplitudes along the first principal component (PC1), indicating continuous conformational heterogeneity in which the helical bundle adopts a range of orientations relative to the catalytic core. To further reduce the impact of this continuous variability present in the full particle dataset, a subset of 109,830 particles with eigenvalues between −3 and 3 along PC1 was selected. These particles were then subjected to an additional round of 3D refinement followed by a second multi-body refinement, resulting in improved signal for the helical bundle of the HAT module.

### Model Building and Validation

The catalytic TIP60 subunit was manually docked into the cryo-EM density using the crystal structure of the TIP60 catalytic core (PDB: 2OU2). The Tudor domain of TIP60 was predicted using AlphaFold^47,48^ and fitted into the corresponding density. EPC1 was modeled based on AlphaFold predictions and manually docked into the cryo-EM map, followed by iterative coordinate adjustment in Coot to optimize agreement with the density^49^. For regions interacting with linker DNA, model interpretation was guided by structural features observed in the yeast NuA4-nucleosome complex^23,24^ together with AlphaFold predicted models^47,48^, which were manually fitted into the corresponding cryo-EM density. The complete atomic model was refined against the cryo-EM map using real-space refinement in Phenix^50^. Multiple sequence alignments were visualized using ESPript^51^. Figures were generated using ChimeraX^52^.

## ACKNOWLEDGMENTS

We thank Dan Toso and Paul Tobias at the Cal-Cryo facility for assistance with high-resolution cryo-EM data collection and computation. Z.Y. is presently an investigator of SUSTech Institute for Biological Electron Microscopy.

## AUTHOR CONTRIBUTIONS

J.C. and E.N. conceptualized and supervised the project. Z.Y. performed the biochemical characterization and structure determination of the TIP60-C HAT–NCP complex. A.J.F.A. assisted with cryo-EM data processing. Z.Y., J.C., and E.N. wrote the manuscript with contributions from all authors.

## DATA AND CODE AVAILABILITY

Structural coordinates of the human TIP60-C HAT-NCP and the corresponding cryo-EM map are being deposited to the Protein Data Bank and the EM Data Bank, respectively. This preprint will be updated with the codes as soon as they are available. Data and materials can be obtained from the corresponding authors upon request.

## EXTENDED DATA FIGURES

**Extended Data Fig. 1.**
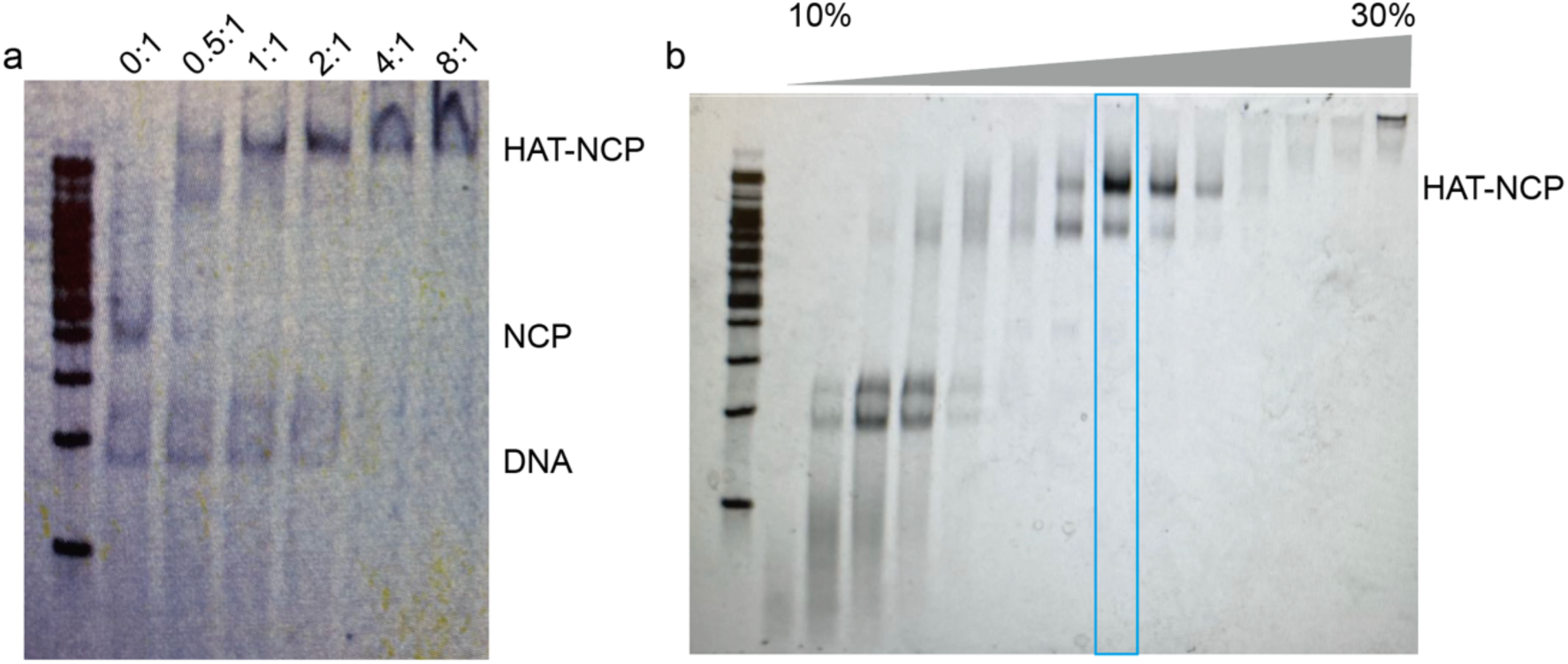
Assembly of the human TIP60-C HAT–nucleosome complex. **a**, EMSA of the TIP60-C HAT module with CMC-modified nucleosomes. At a HAT:nucleosome molar ratio of 4:1, the nucleosome is fully shifted. **b**, GraFix stabilization of the HAT–nucleosome complex. Fractions collected from top (10% glycerol) to bottom (30% glycerol) are shown. The fraction enriched for the HAT–NCP complex (highlighted by the blue box) was used for structural analysis.

**Extended Data Fig. 2.**
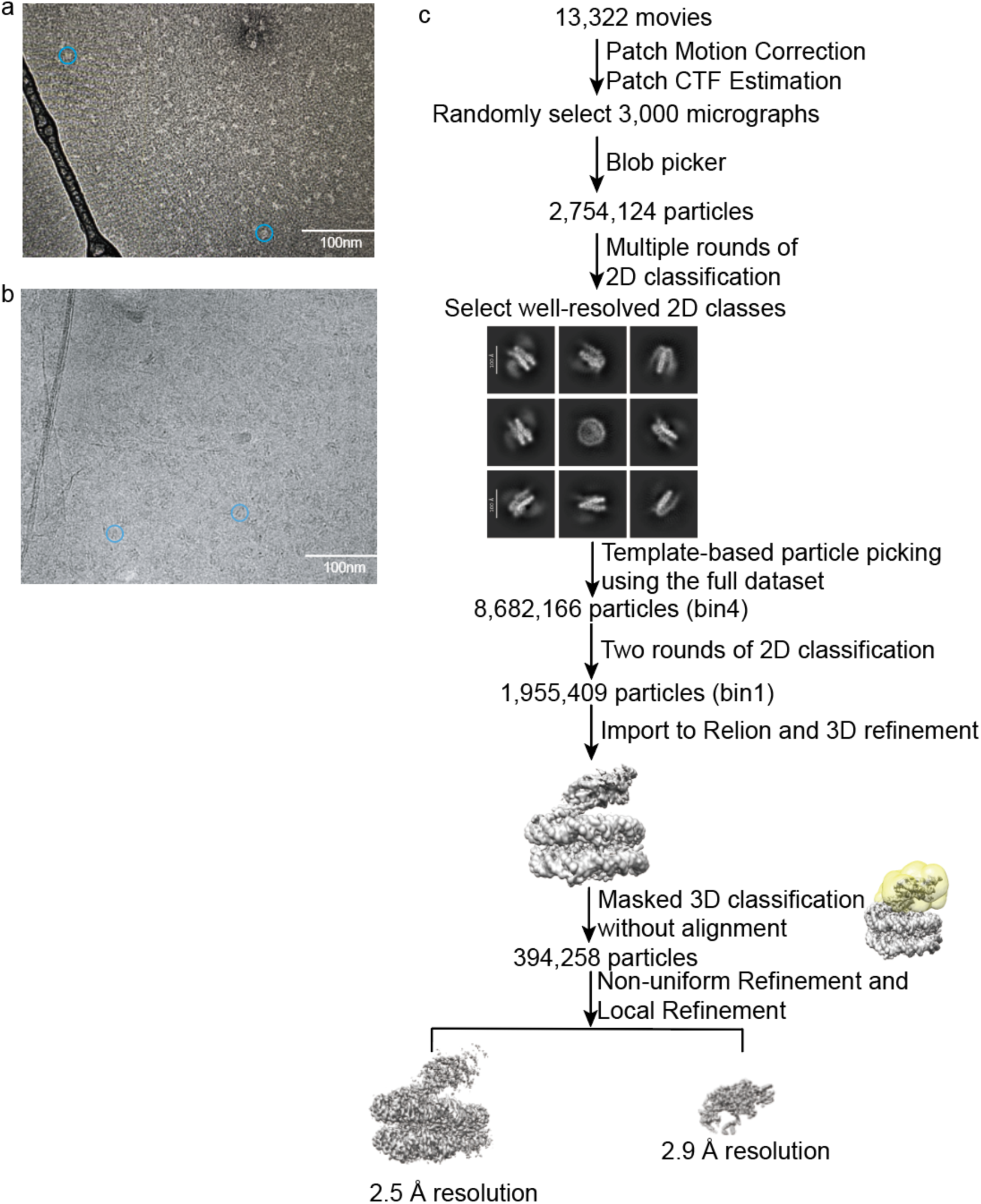
Cryo-EM analysis of the TIP60-C HAT–nucleosome complex. **a**, Example of negative-stain EM micrograph with a couple of examples of target particles indicated by blue circles. **b**, Example of cryo-EM micrograph with a couple of target particles indicated by blue circles. **c**, Workflow for the cryo-EM data processing of the TIP60-C HAT–nucleosome complex.

**Extended Data Fig. 3.**
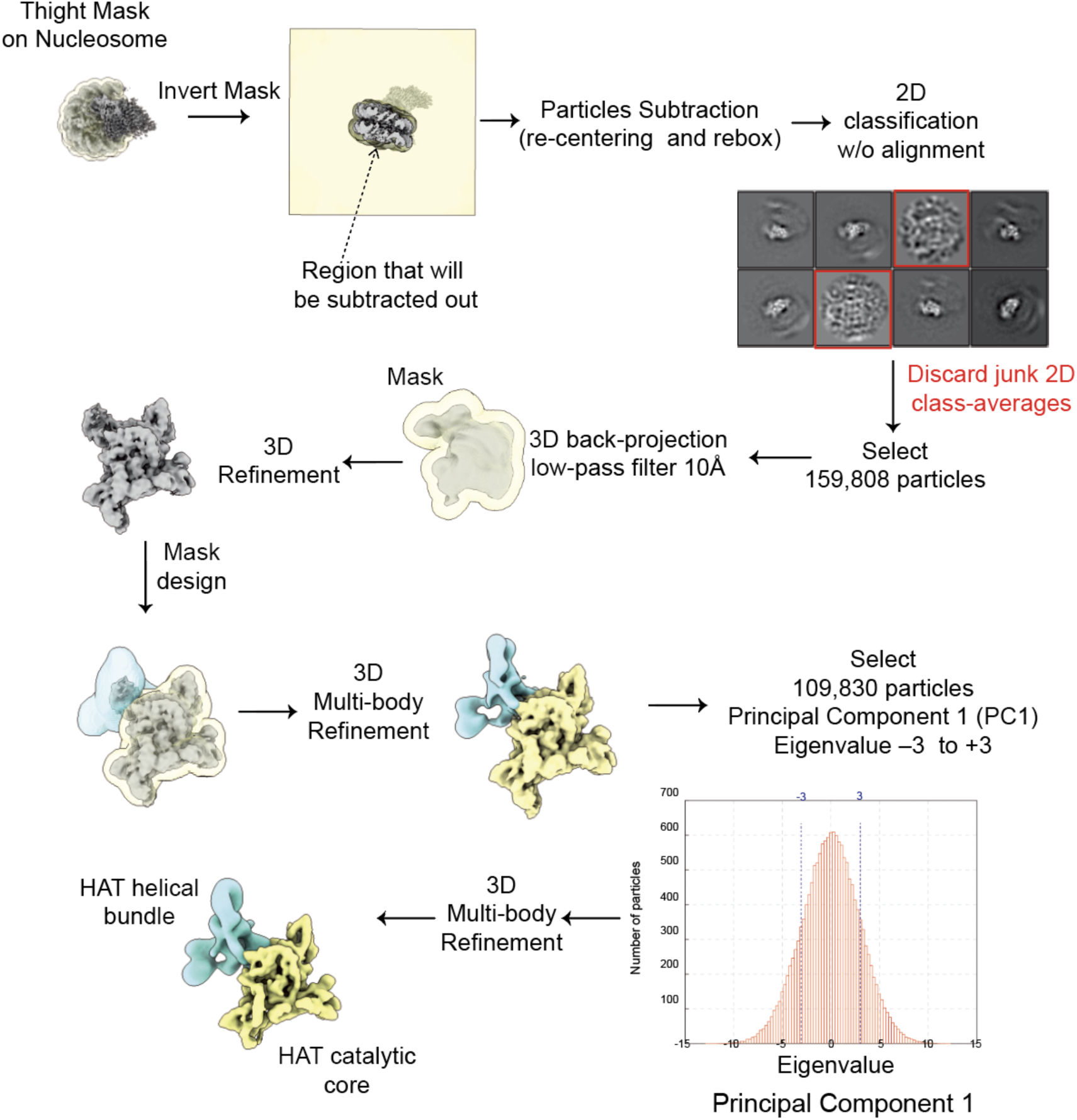
Cryo-EM analysis of the TIP60-C HAT–nucleosome complex used to improve the density of the helical bundle.

**Extended Data Fig. 4.**
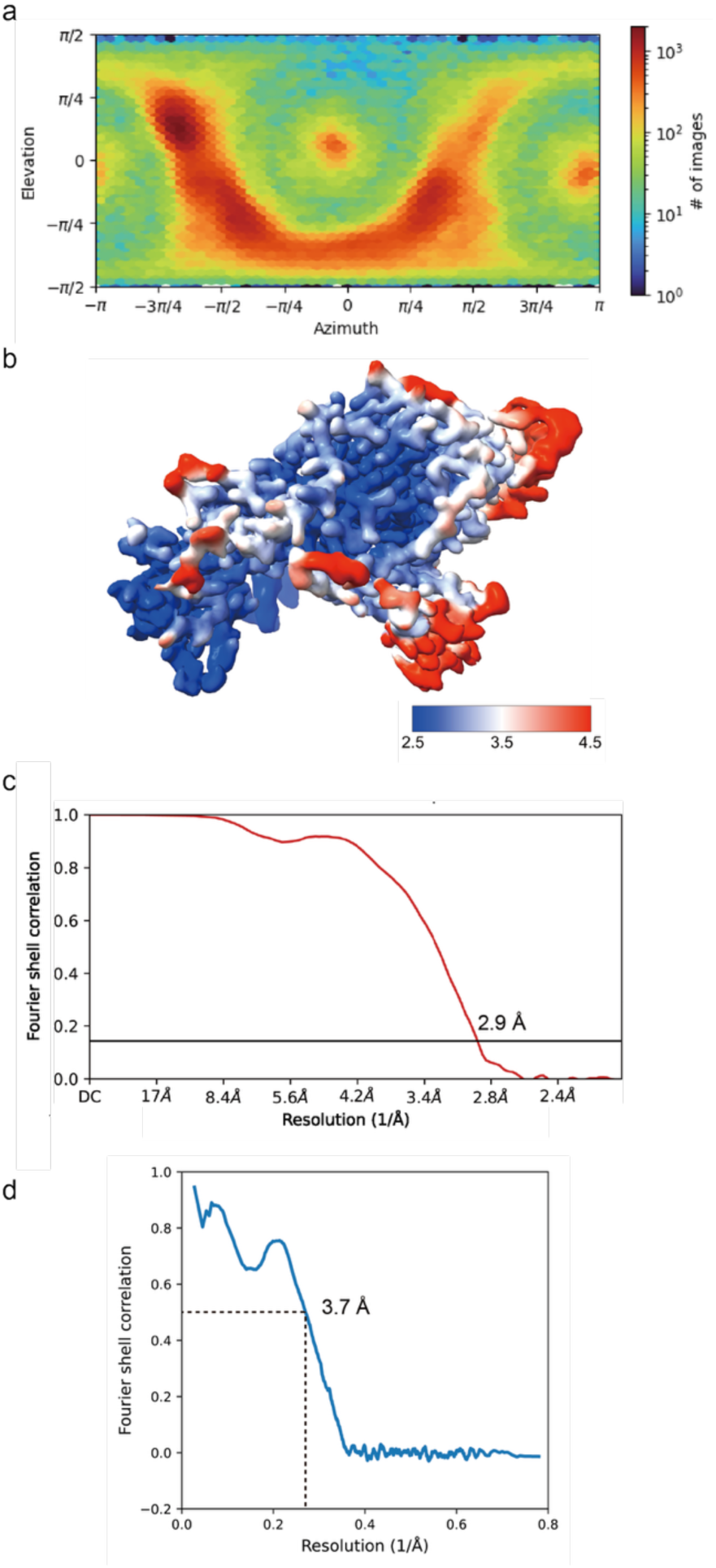
Map quality analysis of the HAT module. **a**, Angular distribution of particles used in the final 3D reconstruction. **b**, Cryo-EM map colored by local resolution. **c**. Gold-standard Fourier shell correlation curve (FSC) of the cryo-EM map (map-map). **d**, FSC curves between the refined atomic model and the cryo-EM map, calculated using Phenix real-space refinement.

**Extended Data Fig. 5.**
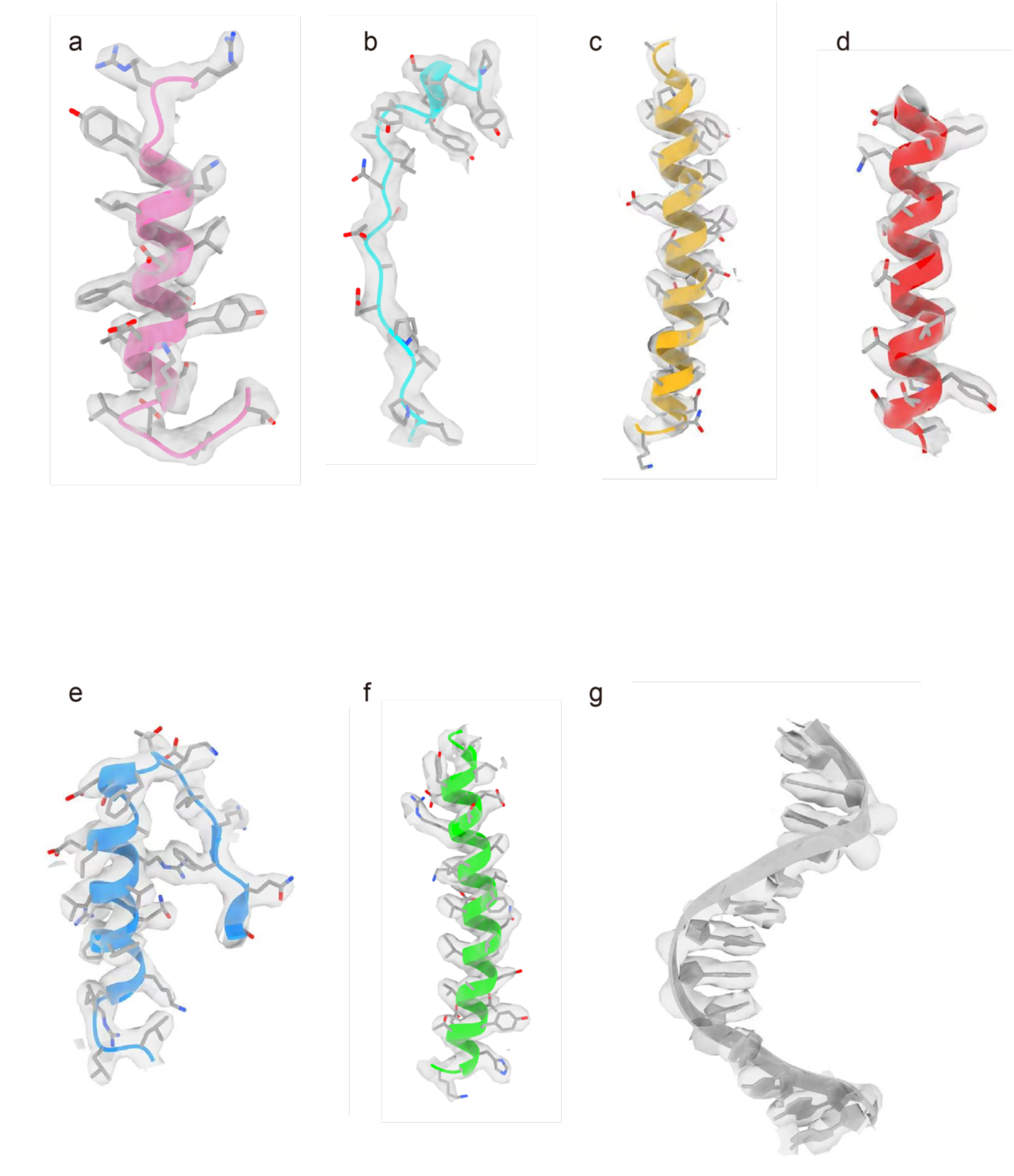
Cryo-EM densities and fitted models for the TIP60-C HAT–nucleosome complex. Representative regions of the cryo-EM density map corresponding to TIP60 residues 378–399 (**a**), EPC1 residues 73–91 (**b**), histone H2A residues 45–74 (**c**), histone H2B residues 102–121 (**d**), histone H3 residues 60–86 (**e**), histone H4 residues 48–77 (**f**), and a single strand segment of nucleosomal DNA for nucleotides 25–37 (**g**).

**Extended Data Fig. 6.**
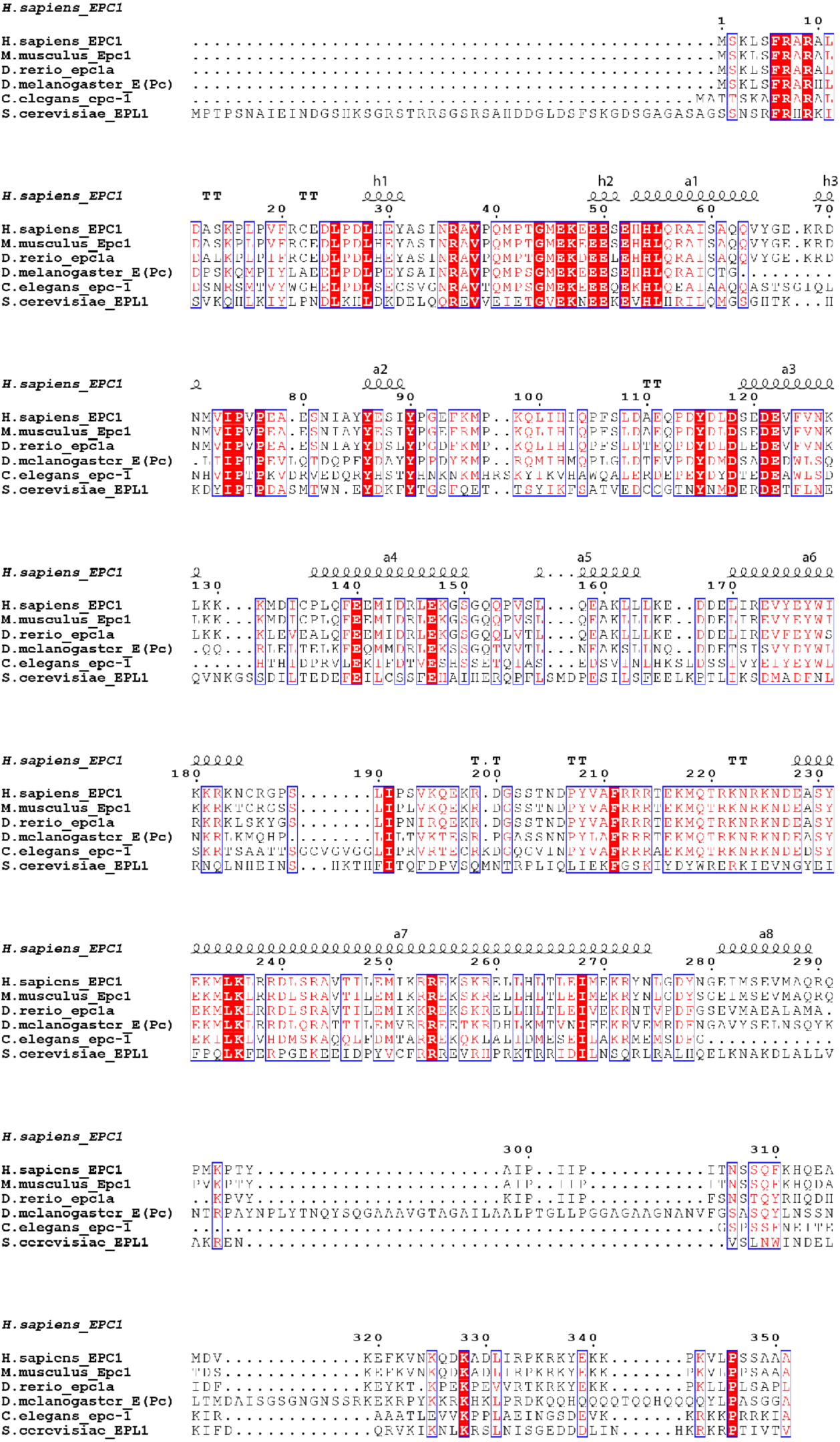
Structure-based sequence alignment of EPC1 across different species.

**Extended Data Fig. 7.**
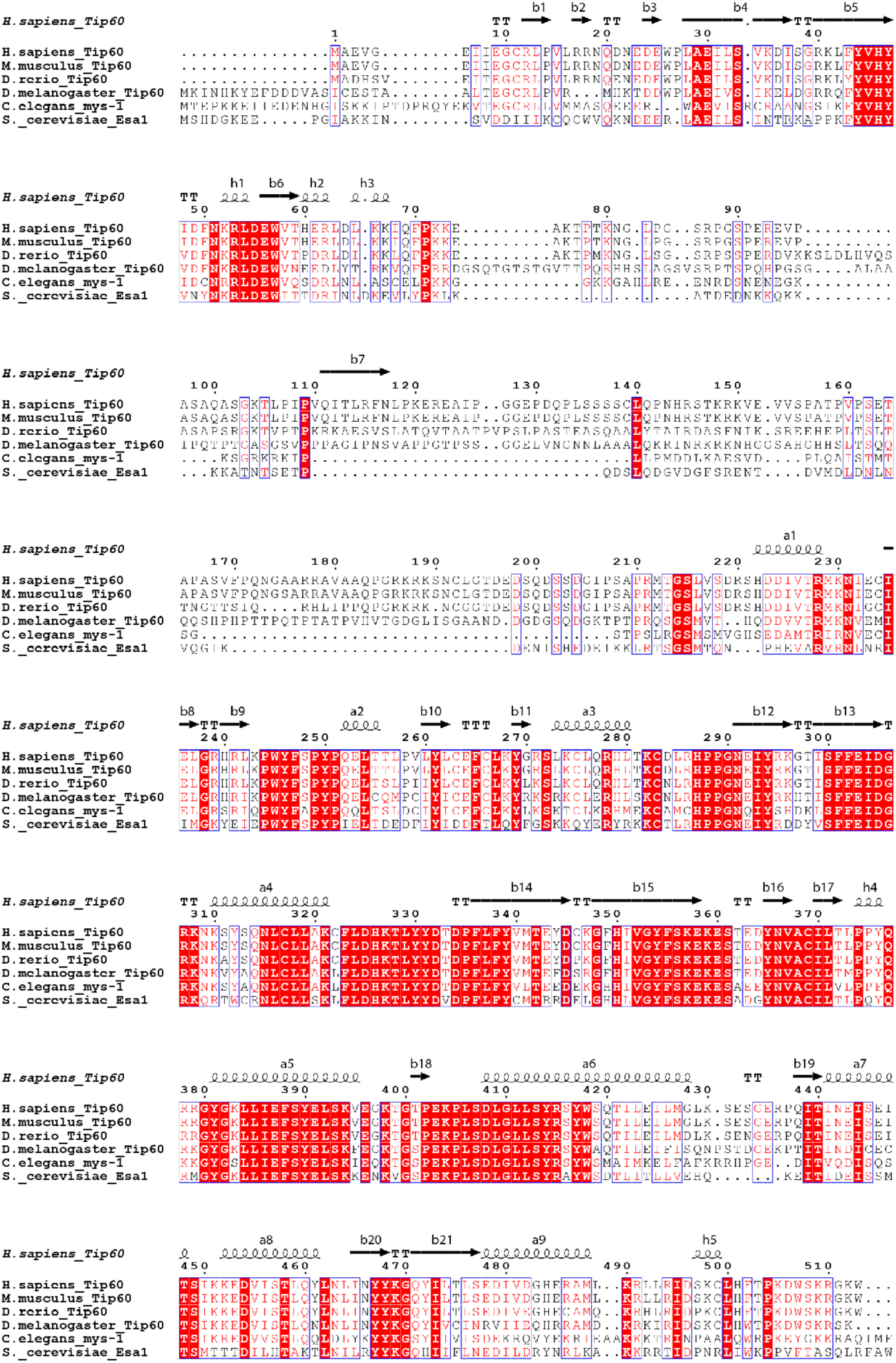
Structure-based sequence alignment of TIP60 across different species.

